# Detection and characterization of a novel bat filovirus (Dehong virus, DEHV) in fruit bats

**DOI:** 10.1101/2023.08.07.552227

**Authors:** Biao He, Tingsong Hu, Xiaomin Yan, Fuqiang Zhang, Changchun Tu

## Abstract

Although bats are natural hosts of most filoviruses (FiVs), with the pathogenic ones frequently causing deadly haemorrhagic fevers in Africa, the viruses are difficult to trace. Recently, multiple divergent FiVs have been uncovered in China, raising concerns about their threat to public health. Here we report the detection of bat-borne FiVs at the interface of orchards between bats and humans, eventually resulting in the discovery of a novel FiV, Dehong virus (DEHV), from *Rousettus leschenaultii* (*R. leschenaultii*) bats. DEHV has the largest genome, and the phylogeny places it between the genera *Dianlovirus* and *Marburgvirus*, suggesting its classification as a prototype of a new genus within the family *Filoviridae*. Our study emphasizes the importance of further understanding the distribution and potential risk of FiVs in the region.

---

Filoviruses (FiVs) are among the most concerning pathogens in the world and are represented by the notorious Ebola and Marburg viruses, which cause highly contagious viral haemorrhagic fevers(1). FiVs, featuring a baculiform morphology, are initially composed of genera *Ebolavirus* and *Marburgvirus* (1). The former consists of very genetically diverse members including four African species, *Zaire ebolavirus* (Ebola virus, EBOV), *Sudan ebolavirus* (Sudan virus, SUDV), *Bundibugyo ebolavirus* (Bundibugyo virus, BDBV), and *Taï Forest ebolavirus* (Taï Forest virus, TAFV), and an Asian species *Reston ebolavirus* (Reston virus, RESTV)(1). Recently, a new ebolavirus species, *Bombali ebolavirus* (Bombali virus, BOMV), has been discovered from bats in Sierra Leone(2). In contrast, the genus *Marburgvirus* is much more restricted in genetic diversity and comprises the unique species *Marburg marburgvirus* with two members, Marburg virus (MARV) and Ravn virus (RAVV)(1).

Recently, two more mammalian FiV genera have been established. The discovery of Lloviu virus (LLOV) in Spanish bats in 2011 formed the genus *Cuevavirus* with it being the only member(3). The latest identification of Měnglà virus (MLAV) in *Rousettus* bats in China has formed another new genus *Dianlovirus* (4, 5). Although diverse mammalian filoviruses have been discovered so far, they show distinct pathogenicity to humans and other animals. EBOV, SUDV, and MARV frequently trigger deadly human haemorrhagic fevers in Africa(6, 7), while the diseases caused by BDBV and TAFV are much rare(6). RESTV is nonlethal to human(1), but fatal to nonhuman primates with several outbreaks in cynomolgus monkeys (*Macaca fascicularis*) in the Philippines(1, 8). LLOV is a possibly deadly agent for bats and presumably responsible for mass mortalities in *Miniopterus schrebrisii* (*M. schrebrisii*) bats in Spain, Portugal, France, and Hungary(3, 9-11), but no human cases have been reported so far. The newly identified MLAV and BOMV have not been reported to be associated with any diseases in humans and other animals.

Bats, especially those in Africa, participate deeply in the circulation of multiple mammalian FiVs. Cell isolation, viral RNA detection, serology survey, and experimental infection have all been accomplished for MARV, providing solid evidence for that African *R. aegyptiacus* bats are natural reservoirs of the virus(12-17). Very recently, LLOV was successfully isolated from the blood of a live *M. schreibersii* bat in Hungary, supporting *M. schreibersii* as the host for LLOV in Europe(10). Members of the genus *Ebolavirus* have not been isolated from bats, although increasingly serological and viral RNA investigations illustrate that a broad range of bat species are involved in the ecological circles of EBOV, SUDV, and BOMV in Africa(2, 18-22). Besides, serological investigations in China(23, 24), Bangladesh(25), the Philippines(26, 27), Singapore(28), and India(29) indicate circulations of RESTV, EBOV-like, and multiple new FiVs in insectivorous and/or fruit bats in Asia, with some further validated by viral RNA detection(26, 30, 31). Of note is southwest Yunnan province, China, where diverse bat FiVs have been uncovered in last ten years. Our viral metagenomic investigation discovered the first divergent FiV DH04 in the lung of a *R. leschenaultii* bat sampled in Yunnan Province(31), which initiated a series of surveys of bat-borne FiVs in the area(4, 24, 30). Those surveys uncovered several divergent FiV genomic fragments in *Rousettus* and *Eonycteris* spp. bats, one of them had its complete genome sequenced, adding the genus *Dianlovirus* to the family(4, 30). In addition, serology surveys of bats across a broad area of south China revealed wide and complex FiV infections(23, 24), suggesting the existence of yet-to-be-discovered new FiV(s) circulating in the area.

Yunnan province is a subtropic area located in southwest of Chinese. It has abundant forests and wildlife resources. The wide plantations of tropical fruits, such as lychee, longan, and jujube, are important sources of livelihood in the region, but also attract a lot of fruit bats to visit at harvest seasons. To prevent those animals from stealing those fruits, orchard owners usually set up nets around orchards to protect their harvest seasons. We initially discovered FiV DH04 from the lung sample of one of 29 *R. leschenaultii* bats trapped in an orchard in this area in 2013(31). Since then, we have been conducting a long-term surveillance of bat FiVs in the area with the focus on several neighboring orchards at harvesting seasons. Finally, a total of 422 fruit bats trapped in orchard protecting nets were collected in five rounds of sampling visits between December 2014 and January 2019, covering *R. leschenaultii* (n = 345), *Cynopeterus sphinx* (n = 65), *Megaerops niphanae* (n = 3), and *Eonecteris spelaea* (n = 9).

We employed a pan-FiV nested RT-PCR(31) to screen all samples and only found the positivity in the lung and the liver of a *R. leschenaultii* bat (sample code Rl133-16) collected in December 2016. The amplicon showed 78.3 - 78.8% nt similarities to the corresponding region of RNA-dependent RNA polymerase gene of FiVs detected in Chinese bats including MLAV, and then 77.4% to that of EBOV, indicating the identification of a novel FiV. We tentatively named it Dehong virus (DEHV) based on the place of its discovery.

We sequenced the complete genome of DEHV by high-throughput sequencing (HTS) and verified the sequence using the traditional Sanger method following RT-PCR amplification. Its complete genome is 20,943 nt in length (Genbank accession number OP924273), the largest genome size in the family *Filoviridae* (Fig. 1a). It has 13 nt-long exactly reverse complementary extreme 3’ and 5’ ends. The extragenic sequences at the 3’ end (leader) is 49 nt long, but much longer at the 5’ end (trailer), having 774 nt (Fig. 1a). The genome contains seven sequentially arranged genes in the order of nucleoprotein (NP)-virion protein (VP) 35-VP40-glycoprotein (GP)-VP30-VP24-polymerase (L), which are separated by four intergenic regions, while the VP35 and VP30 are partially overlapped by VP40 and VP24, respectively (Fig. 1a). The genome structure is more similar to that of MARV, RAVV, and MLAV than to that of the remaining FiVs. These genes are delineated by the conserved transcriptional signals, which are the same as those in MARV and RAVV. The transcriptional start and stop signal for DEHV are 3’-CUWCUURUAAUU-5’ and 3’-UAAUUC(U)5-5’, respectively (Fig. 1a). The GP of LLOV and ebolavirus species is encoded in two open reading frames (ORFs) and expressed through transcriptional editing, while like marburgviruses and MLAV, the GP of DEHV is encoded in a single ORF (Fig. 1a).

**Figure 1.**
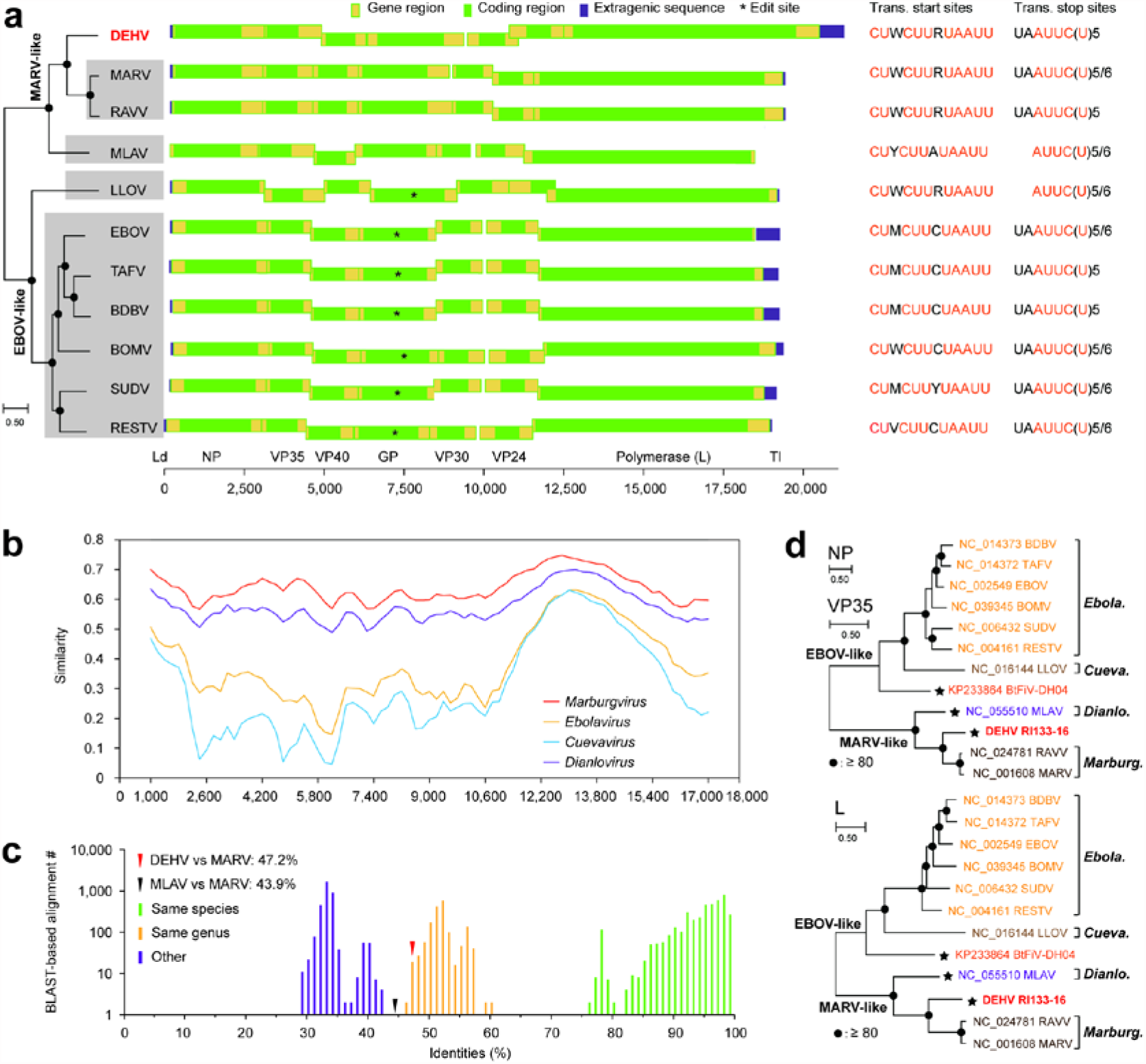
Genomic and phylogenetic characterization of DEHV. (a) Genomic structure comparison of DEHV with other FiVs. The phylogenetic tree is based on the complete genome sequences using maximum-likelihood (ML) method. The node with filled circle indicates that the divergence is supported by 75% bootstrap replicates. (b) The similarity profiles of DEHV with the consensus sequences of the genera *Marburgvirus, Ebolavirus, Cuevavirus*, and *Dianlovirus*. (c) Taxonomy assignment of DEHV based on ICTV proposed BLAST-based alignment comparison. (d) The ML phylogenies of DEHV with other representatives of mammalian FiVs based on NP, VP35, and partial L aa sequences. The phylogenetic trees of NP and VP35 are highly identical, hence are represented using a consensus tree. The node marked using filled circles indicates it is supported by 75% or 80% bootstrap replicates.

The global alignment based on the complete genomes revealed that DEHV has the highest ∼60.0% nt identity to MARV and RAVV, followed by 53.9% to MLAV, but distantly related to ebolaviruses (39.9-41.0%) and LLOV (37.2%). DEHV shows similar sequence similarity profile across the entire genome to currently known mammalian FiVs, i.e., higher similarity at the central region of the NPs and the upstream region of the L gene (Fig. 1b). The similarity profile did not find any intersection between different FiVs (Fig. 1b), suggesting long-term independent evolution among FiV lineages. No evidence statistically significant for historical recombination events among filoviruses was found by using PhiTest (p = 1.0) and RDP4. The International Committee on Taxonomy of Viruses (ICTV) proposed an algorithm for filovirus classification based on genomic sequence information and Pairwise Sequence Comparison (PASC)-derived sequence demarcation criteria(32). Genomic sequences of different filovirus genera should differ from each other by ≤ 45% identities based on the BLAST alignment(32). By this method, DHEV showed the highest identity with MARVs (46.3 - 47.2%), slightly higher than the approved genus criteria (Fig. 1c). Viruses with close genetic relationships usually have compatible nucleotide frequency. The test of the heterogeneity of nucleotide frequencies among different FiVs showed that DEHV had a significantly different nucleotide frequency pattern (Chi-square test, 𝒳^2^ = 1577.3, P < 0.0001) as compared to other FiVs. It has the highest AU content (68.2%), while such contents for marburgviruses, MLAV, LLOV, and ebolaviruses are 61.7-62.2%, 63.5%, 54.0%, and 54.5-59.4%, respectively.

Phylogenetic analyses revealed that the seven deduced ORF amino acid (aa) sequences, as well as the complete genomes, of these mammalian FiVs show highly identical phylogenetic topology (Figs. 1a, d). FiV DH04 was also included in the phylogenetic analyses of NP, VP35, and L (Fig. 1d). These results showed that FiVs are polyphyletic and are clearly segregated into two major phylogroups named respectively as EBOV-like and MARV-like phylogroups, which were interestingly corresponded to whether they have edit site in GP gene (Fig. 1a). The EBOV-like phylogroup consists of DH04, LLOV, and six ebolaviruses, while the MARV-like one comprises DEHV, MLAV, and two marburgviruses (Fig. 1d). DEHV is placed between MLAV and marburgviruses, but relatively closer to marburgviruses (Fig. 1d).

To understand the dynamics of natural DHEV circulation in the region, a specific RT-PCR method modified from the pan-FiV nested RT-PCR using primers designed based on DEHV genome sequence was used, along with a qRT-PCR method, to re-screen the 422 bats. It further confirmed the positivity of the lung and liver of bat Rl133-16 with RNA loads of 8.0 × 10^4^ and 2.6 × 10^5^ genomic copies/0.1 g, respectively. In addition, we identified one more positive lung sample of *R. leschenaultii* bat Rl81-14 collected in December 2014 with an RNA load of 1.7 × 10^4^ genomic copies/0.1g. Further sequence comparison showed that the 249 nt-long FiV contig previously identified by the metatranscriptomic sequencing of the pooled sample of 57 *R. leschenaultii* bats collected in January 2019 in the same region(24) is 100% identical to L gene of DEHV, but unfortunately the above two specific RT-PCR methods failed to detect any positivity by re-screening those 57 bats, indicating a very low viral load in these samples. The continuous detection of DEHV in *R. leschenaultii* bats in 2014, 2016, and 2019 in these orchards suggested a natural circulation of DEHV in the region.

Interface for contact between human and wild animals is a key factor mediating spillover of wildlife-borne viruses to cause public health risk. Orchards in bat dwelling areas are such interfaces where fruit bats forage at night in various fruit trees at harvesting seasons. Since the first discovery of FiV DH04 in a *R. leschenaultii* bat collected in an orchard in Yunnan Province, 2013(31), we have been conducting a long-term surveillance to track bat-borne FiVs in the region in order to assess the risk of their potential spillover. Here we report the milestone results that reflect how we tracked bat-borne FiVs during the last ten years through continuous sampling and detection of fruit bats, eventually resulting in the identification and isolation of DEHV from *R. leschenaultii* bats. We conducted 5 rounds of samplings of fruit bats trapped in the orchards in 2014 (Dec), 2016 (Jul-Aug and Dec), 2017 (Dec), and 2019 (Jan), and pan-FiV RT-PCR detected only one positive bat (*R. leschenaultii*) from samples of 2016 (leading to the discovery of DEHV), while all other samples from 2014 and 2019 were negative. To investigate the natural circulation of FiVs in bats in the region and other south provinces, we analyzed FiV seroprevalence of 689 archived serum samples of 16 bat species collected between 2012 and 2016 in previous study(24), which showed 36.3% overall seropositive rate against multiple FiVs, and identified two unknown FiV reads by additional viromic sequencing from the pooled samples of 57 *R. leschenaultii* bats collected in December, 2019, in these orchards(24). The two reads were re-analyzed upon the discovery of DEHV, and surprisingly showed 100% identity to the corresponding regions of L gene of DEHV, indicating that the FiV detected in the previous study(24) was DEHV. Therefore, we decided to re-screen all samples of those 422 bats collected between 2014-2019 by DEHV-specific RT-PCR and qRT-PCR. Unfortunately, no positive detection was from any of those 57 *R. leschenaultii* bats, but surprisingly we identified one more positive *R. leschenaultii* bat from 120 samples of 2014. In other Asian countries serological investigations revealed high FiV seroprevalences in several bat species in the Philippines(26, 27), Singapore(28), Bangladesh(25), and Indian(29), but the virus has not been identified except for RESTV detected from bats (mainly from *M. schreibersi*) in the Philippines(26). In China, several frugivorous and insectivorous bat species also have high seroprevalence mainly in the south provinces(23, 24), with FiV RNA detected only in *Rousettus* spp. and *E. spelaea*(30, 31). *R. leschenaultii* and *E. spelaea* bats are widely distributed with dense populations throughout south China, southeast Asia, and/or the entire Indian subcontinent(33). Of these bats, *R. leschenaultii* has the highest frequency of virus detection, which had 4 times of FiV-positive detections in 2013, 2014, 2016 and 2019 in our previous(24, 31) and present studies. In addition, another team detected different FiVs from *Rousettus* spp. and *E. spelaea* bats collected in 2009 and 2015 from Jinghong and Mengla of Xishuangbanna Prefecture(30) (Fig. 1a), leading to the identification of MLAV from one *Rousettus* sp. bat(4). Our serological investigation showed 60.8% (87/143) seroprevalence against FiV in *R. leschenaultii* bats in Xishuangbanna Prefecture(24). Serological study on bats collected between 2006-2009 in this region by Yuan *et al*., showed that the most significant prevalence of ebolavirus antibody was found among *R. leschenaultii, Pipistrellus pipistrellus* and *Myotis* sp.(23). These results indicate that *R. leschenaultii* is a major reservoir species harboring multiple bat FiVs in Yunnan province.

Based on our phylogeny, mammalian FiVs could be segregated into MARV- and EBOV-like phylogroups (Fig. 1a). All members in the former (DEHV, MARV, RAVV, and MLAV) are monophyletic, harbored by frugivorous *Rousettus* bats and have the same encoding strategy of GP (Fig. 1a), suggesting that they are progenies of a consensus ancestor virus infecting *Rousettus* bats. Notably, frugivorous bats in Yunnan province harbor more diverse FiVs, including DH04, MLAV, DEHV, and even a number of yet-to-be-identified FiVs(30), and these viruses are polyphyletic with DH04 and MLAV being basal in the phylogeny (Fig. 1d). Therefore, these MARV-like FiVs likely originated from and have circulated in Asian *Rousettus* bats for a long time. Considering the distribution of FiV-positive bats along border areas and the long distance *R. leschenaultii* is able to migrate(33), the circulation area of DEHV is wider than just the place it was discovered. Additionally, DH04, MLAV and other yet-to-be-identified FiVs were detected in Jinghong and Mengla(4, 30), thereby a natural circulation sphere of diverse FiVs covering Yunnan province and bordering Southeast Asia has been proposed, where diverse MARV-like FiVs are harbored by *R. leschenaultii* or even other bat species. This region is another FiV hot spot in addition to Central Africa, West Africa, and South-Central Europe. The finding highlights the importance of further understanding the pathogen ecology of bat-borne FiVs in the future.

There are biological traits shared by DEHV and the other MARV-like FiVs, but DEHV is phylogenetically distinct. Its genome is the largest, at least 1,800 nt longer, and shows different base frequency with much higher AU content as compared to other MARV-like FiVs. In addition, the PASC shows that the highest BLAST-based identity of DEHV to MARV is very close to the genus demarcation criteria. Therefore, it is acceptable to classify DEHV as a member of a new genus within the family *Filoviridae*. As per *Dianlovirus*(4), we propose the genus name as *Delovirus*, reflecting a new fi*lovirus* discovered in *De*hong Prefecture of Yunnan province, and DEHV is the prototype of the species *Dehong Delovirus* within the genus.

## Materials and Methods

Between December 2014 and January 2019, we collected bats trapped in protective nets that were set up at several adjacent orchards planting lychees, longans, and jujubes, which were very close to human residences. These collections were conducted at five fruit harvest seasons with each spanning 6-20 days. We inspected these nets every morning and picked trapped bats from nets. Most bats were adult, but their exact ages and genders were not recorded at sampling. Majority of these bats were dead at collection, and they were immediately subjected to necropsy at the local laboratory to sample their brains, lungs, livers, kidneys, spleens, and recta, and sera when possible. All samples were cryo-transported to our laboratory and stored at -80 °C. The species was morphologically identified and further confirmed by sequencing the mitochondrial cytochrome b (*Cyt b*) gene (34).

### Virus detection

All samples were subjected to the pan-FiV detection as per our published method (31), which uses nested degenerate primer pairs focusing on the most conserved region of L gene of all known mammalian filoviruses and was successfully employed to detect diverse bat FiVs (30). Briefly, about 0.1g of each tissue was subjected to total RNA extraction using a QIAamp RNeasy Mini Kit (Qiagen, Hilden, Germany). The cDNA was synthesized for PCR reaction using a PCR master mix (Tiangen, Beijing, China). The specific amplicons were cloned into pMD18T vectors (TaKaRa, Dalian, China), and 10 clones of each amplicon were randomly picked for sequencing by the Sanger method on an ABI 3730xl DNA sequencer (ComateBio, Changchun, China).

To specifically detect DEHV, above degenerate primers were replaced by specific ones based on the genome sequence of DHEV. Besides, DHEV-specific qRT-PCR was established to quantify the virus load of positive samples with the primer and probe sequences being: DHEV-GP-QF1 (5’-ACTGAACCAGGCGAGAAGAAA-3’), DHEV-GP-QR1 (5’-ACCAGGACCAAAAAAGGGAAT-3’), and DHEV-GP-probe (HEX5’-CCATCTCGTCCTCTTGAATGCTCCATA-3’MGB). The plasmid bearing the amplicon was constructed to generate the standard curve following serial 10-fold dilutions. These positive tissues were accurately weighed and completely homogenized for total RNA extraction. The qRT-PCR program was performed using the MonAmp Taqman qPCR mix (Monad, Wuhan, China) using the following program: 15 s at 95 °C, then 40 cycles of 5 s at 95 °C, 30 s at 60 °C.

### High-throughput sequencing

To obtain the complete genomic sequence of DEHV, RNA was extracted using phenol-chloroform and subjected to ribosomal RNA removal using a NEBNext^®^ rRNA Depletion Kit (Human/Mouse/Rat) (NEB, Ipswich, MA). A RNAseq library was prepared by using a NEBNext^®^ Ultra directional RNA library prep kit (NEB) and then sequenced on an Illumina NovaSeq 6000 sequencer. About six gigabase raw data were generated. These reads were quality checked using fastp version 0.20.0, and the resultants were *de novo* assembled using SPAdes genome assembler version 3.14.1 with meta mode. Contigs of ≥ 1 kb were used for ORF prediction using prodigal version 2.6.3 with the translation table 1 and meta mode. The derived amino acid sequences of ≥ 50 aa were queried against the Eukaryotic Viral Reference Database (EVRD)-aa version 1.0(35) using diamond blastp version 0.9.35 with e-value cutoff 1e-5. This search found a 21,937 nt-long contig corresponding to a new FiV. We performed self-to-self alignment using the reverse complementary sequences to identify the extreme 3’ and 5’ ends, and further validated them using RT-PCR and Sanger sequencing. To check the quality of assembly, we mapped these reads back to the complete sequence using bowtie2 version 2.4.1, and calculated the sequencing coverage using samtools version 1.10.

### Sequence comparison and phylogenetic analysis

The representative sequences of mammalian FiVs were retrieved from RefSeq. They are EBOV (NC_002549), BDBV (NC_014373), TAFV (NC_014372), BOMV (NC_039345), SUDV (NC_006432), RESTV (NC_004161), LLOV (NC_016144), MLAV (NC_055510), RAVV (NC_024781), and MARV (NC_001608). The partial cds of BtFiV DH04 was also withdrawn from GenBank. These sequences were all used in the sequence and phylogenetic analysis with DEHV. The ORFs of DEHV were *de novo* predicted using ORF Finder (www.ncbi.nlm.nih.gov/orffinder) and further compared with those of other known representative sequences. The locations of transcriptional start and stop signals of DEHV were determined by searching the consensus motif of the other FiVs. Base composition bias refers to uneven distribution and composition of four bases in a genome. Usually, base composition is stationary among genomes or genic homologs of close taxa. The base compositional heterogeneity of DHEV with other filoviruses was tested using DAMBE version 7.3.11(36). The pooled sequence set (i.e., all complete genomes of FiVs) was first subjected to Chi-square test, if the set did not show homogeneity of base frequencies across taxa (P < 0.05), the sequences were then pairwise-tested. To assess the global sequence identity of DHEV with other FiVs, their complete genomes were aligned using mafft version 7.470 with e-ins-i method, and the pairwise identities were calculated using MegAlign module within the Lasergene suite version 7.1.0. PASC, the BLAST-based alignment web tool proposed by ICTV, was used to classify a sequence within the family *Filoviridae*(32). Accordingly, we used the strategy in the taxonomic assignment of DEHV.

Mutation and recombination are two important natural drivers of shaping genetic diversity of non-segmented RNA viruses. In terms of negative-sense RNA viruses, homologous recombination occurs in mumps virus and Newcastle disease virus within the family *Paramyxoviridae*, and rabies virus and spring viraemia of carp virus within the family *Rhabdovirdae*, though considerably lower than rates of mutation(37-40). Among rabies viruses, recombination has been detected between different lineages and presumably promoted the adaptation of the viruses to carnivores by host shift from bats(40). To detect the historical recombination possibilities among filoviruses, their extent of recombination was first detected using PhiTest with window size of 100, which is a robust statistical method for detecting the presence of recombination(41). The phi test did not find statistically significant evidence for recombination (p = 1.0). The detection was then confirmed by a series of prediction implemented in RDP4(42), in which nine methods (i.e., RDP, GENECONV, BootScan, MaxChi, Chimaera, SiSican, 3Seq, LARD and PhyIPro) were employed and if a recombinational prediction was simultaneously supported by at least three methods with MC-corrected probability ≤ 1e-3, it was considered significant.

The phylogenies of seven ORFs in aa and the complete genomes in nt were inferred using MEGA version X. The alignments were built using mafft, and their hypervariable regions were automatically trimmed by trimAL version 1.2 with default settings. The best substitution models were assessed using ModelFinder version 1.6.8 with the Akaike information criterion. The maximum likelihood trees were reconstructed using the Subtree-Pruning-Regrafting fast method. The phylogenies were tested using the bootstrap method under 1000 replicates.

## Notes

### Competing Interest Statement

The authors have declared no competing interest.

